# Predicting lead hotspots in urban stormwater ponds across the Twin Cities Metro

**DOI:** 10.1101/2025.11.13.688297

**Authors:** L. Pollack, C. Zweifel, B. Janke, T. Ting, Z. Culshaw-Klein, J.C. Finlay, C. Santelli, E. Snell-Rood

## Abstract

Stormwater ponds play an important role in urban food webs. Critically these same ponds could also serve as pollution hotspots since stormwater ponds can act as local concentrators of urban runoff. One such contaminant of concern is lead, which remains a significant issue for human and ecosystem health in the United States despite regulatory bans on its use in paint and gasoline imposed in the 1970s. Despite high levels of lead in some urban stormwater, little is known about the distribution of lead in urban aquatic ecosystems. Therefore, we characterized lead within the sediment, water column, and surrounding soil of stormwater ponds across the Minneapolis-Saint Paul Metro Area. We hypothesized that lead would be highest in ponds that receive runoff from landscapes with older construction (i.e., legacy leaded paint and gasoline), have high traffic volume (i.e., legacy leaded gasoline), and areas with low impervious surface cover (i.e., increased mobilization of contaminated soil). Moreover, we hypothesized that stormwater ponds capture lead within sediments, with more dissolved lead at the bottom of the water column, where it would interact with lead containing sediments. Across ponds, we found that age of parcel development where the pond was located was the strongest predictor of surface sediment lead content. Within pond sediments, we found that lead concentrations increased with depth below the sediment surface, which is unsurprising since depth is likely correlated with time. The strongest predictor of surface water lead concentration was the strength of pond stratification, while the strongest predictor of bottom water lead concentration was pond duckweed cover and water conductivity. Water column oxygen concentrations varied across ponds yet were not important in determining dissolved lead within the water column. Importantly, lead within pond water remained quite low despite elevated sediment lead levels. These findings confirm that stormwater ponds can act as one source of environmental lead remediation by capturing lead within sediments under a wide range of environmental conditions. Our results suggest relatively low lead release from ponds to downstream areas, indicating that ponds generally serve as sinks, not sources within the urban lead cycle.

## Introduction

Stormwater ponds are fundamental components of anthropogenic ecosystems, acting as critical “control points” for the cycling of elements, including contaminants such as heavy metals. Indeed, there are more stormwater ponds than natural lakes in the United States, making them an important part of modern ecosystems (Olsen 2017, Sinclair et al. 2020, U.S. EPA 2024). A crucial component of stormwater management infrastructure, stormwater ponds help ease downstream flooding and erosion by slowing the flow of water during storm events (MPCA 2025). Critically, stormwater ponds also settle out sediments, particulate matter, nutrients, and other contaminants (Hvitved-Jacobsen et al. 2010, Adhikari et al. 2023), naturally filtering stormwater. For example, when contaminated waters move through stormwater ponds, contaminants like heavy metals are less likely to leach into groundwater or move onto other waterways (Rycewicz-Borecki et al. 2016, Yang et al. 2020). As a result, these systems not only manage hydrological extremes but are also a location of contaminant hyperaccumulation. Their function can influence the long-term fate of pollutants and nutrients within city landscapes, making them potential hotspots for ecological interactions between contaminants and urban biodiversity.

Importantly, stormwater ponds also serve as constructed ecosystems (Sinclair et al. 2020), acting as important reservoirs for urban vegetation and wildlife (e.g., amphibians, Le Viol et al. 2012, Holtmann et al. 2017) and providing habitat for diverse freshwater taxa (Hassall and Anderson 2015, Oertli and Parris 2019, Ferzoco and McCauley 2024). Thus, although pollutants are in theory sequestered within stormwater ponds, interactions within pond food webs could move contaminants into urban freshwater ecosystems and even back into the terrestrial through semi-aquatic vectors like aquatic insects or mallards. Similarly, biogeochemical processes could lead to the unintended release of pollutants from stormwater ponds (Taguchi et al. 2020). More work is needed to confirm if ponds are truly capturing certain contaminants or creating another hub within the urban ecosystem for pollution redistribution. To best understand where and how contaminants are moving into urban ecosystems and the potential impact on aquatic life, we first need to understand where it is accumulating across the landscape of urban stormwater ponds and within the ponds themselves to best locate potential environmental exposure points.

Lead is a pervasive environmental toxin with well-documented risks to both ecosystem and human health, even at low levels of exposure (Mielke 1999, Tong et al. 2000, Kumar et al. 2020, Swaringen et al. 2022). Lead exposure has documented negative impacts on plant and animal communities, ultimately impacting ecosystem services. For example, lead inhibits plant growth and is associated with less abundant and diverse pollinator communities (Moroń et al. 2012, Küpper 2017, Mitra et al. 2019, Zulfiqar et al. 2019). Moreover, lead biomagnifies through food webs, with research documenting neurological and reproductive impacts in birds and aquatic organisms (Scheuhammer 1987, Kane et al. 2005, Baos et al. 2006, De Vries et al. 2007, Bichet et al. 2013, Botté et al. 2022). Unlike organic pollutants, lead is a non-degradable heavy metal that accumulates in soils, dusts, and sediments, making mitigation methods difficult (Dagdag et al. 2023). Lead’s persistence means it remains present in older housing, soil, and aging infrastructure, and can be reintroduced into the air and dust (Levin et al. 2021). As a result, lead is still detectable in human blood and remains a major public health issue (McFarland et al. 2022, Centers for Disease Control and Prevention 2025). In human children, lead exposure is strongly associated with effects that range from decreases in hearing and postnatal growth to increased behavioral problems (Canfield 2003). Adults are also at risk, with increased rates of hypertension, reproductive issues, and cognitive decline later in life associated with lead exposure (Mason et al. 2014, Gambelunghe et al. 2016, Maloney et al. 2018, Antoniadou et al. 2020, Tsoi et al. 2021). These widespread and persistent effects highlight the urgent need to minimize lead contamination to protect both the environment and public health. This study seeks to develop a predictive framework for understanding how stormwater ponds mediate lead movement through urban ecosystems.

At the landscape level, urban soil contamination is likely the most important factor impacting lead accumulation within pond sediment. Studies suggest that soils are a substantial reservoir of environmental lead contamination (Laidlaw and Filippelli 2008, Mielke et al. 2011, Datko-Williams et al. 2014, Laidlaw et al. 2017). Unlike air or water, where lead concentrations can dissipate or be diluted over time, lead persists in soils for decades, making them active repositories and long-term sources of exposure through processes like the resuspension of contaminated dust (Filippelli and Laidlaw 2010, Strosnider et al. 2017). It is well documented that lead moves from urban soils into bodies of water through stormwater (Pitt et al. 1999, Rycewicz-Borecki et al. 2016, Yang et al. 2020). Thus, soil contamination makes its way into freshwater ecosystems when it is captured as sediments in ponds and lakes. The distribution of lead in urban soils is highly heterogeneous, with the highest concentrations typically correlated with vehicle traffic and land use patterns, reflecting the past deposition of paint and gasoline (Mielke et al. 2011, Taylor et al. 2013, Filippelli et al. 2018, Wade et al. 2021). Moreover, green space and impervious surface cover could impact the movement of contaminated soil runoff into ponds by influencing the accumulations, heterogeneity, and erosion of surrounding soil (Chin 2006, Obropta and Del Monaco 2018). In theory, ponds surrounded by more green space would have potentially less aging nearby infrastructure and less runoff with contaminated road dust. However, soil that is buried beneath asphalt and other structures has been shown to be more isolated and thus less likely to be mobilized by stormwater (e.g., abatement strategies reviewed in U.S. EPA and Batelle Memorial Institute 1998, Laidlaw et al. 2017). Thus, increased impervious surface cover can decrease potential transport pathways from urban soils into stormwater ponds.

When considering how lead is then exposed to plants and animals living within stormwater ponds, it is crucial to understand how much lead is dissolved within pond water versus sequestered in less bioavailable forms within the sediment. Critically, lead transforms into different molecular compounds once added to soils and sediments, which can influence its solubility, mobility, and ultimately its toxicity (Manceau et al. 2002, Kushwaha et al. 2018, Haque et al. 2021). Internal features of these ponds likely drive the physical and chemical conditions necessary for dissolving lead or keeping it in less soluble forms. Lead cations (Pb^2+^), when dissolved in water, readily adsorb onto solid surfaces (Lee et al. 2013) and complex with organic matter (Landrot and Khaokaew 2018, Dewey et al. 2021). Dissolved oxygen could create a chemical environment that is more or less conducive to reactions that dissolve lead, but this depends on the ways in which lead is bound as sediment. For example, oxygen rich water could lead to an increase in metals complexed with organic matter due to sediment redox conditions (Miao et al. 2006), but this also depends on the other sediment minerals present (Algeo and Maynard 2008). Furthermore, under anoxic conditions, iron oxides may undergo reduction, releasing bound lead back into the water, where is can subsequently be readsorbed onto organic matter (Cornu et al. 2009, Weber et al. 2009). Critically, many pond features impact anoxic conditions within these ponds, influenced by depth, stratification, vegetative cover, and wind access (Ahmed et al. 2022). Moreover, many urban ponds, especially in snowy climates, can have high salt inputs from frequent winter road salt application (Snodgrass et al. 2017, Janke and Finlay 2025). This salinity would be expected to increase the solubility of lead (Acosta et al. 2011). This is because chloride can complex lead in the aqueous phase, which increases the amount of lead ions the water can contain (Santucci and Scully 2020). Furthermore, calcium rich waters could lead to higher rates of cation exchange (Zhao et al. 2021, Tipper et al. 2021, Miranda et al. 2022), potentially removing some lead from binding sites in the sediment under certain environmental conditions.

The goal of this study is to improve our understanding of where lead is accumulating within the urban stormwater network and improve predictions about where to focus future remediation work. We aimed to determine how land use past and present influence the heavy metal content of stormwater ponds, and where specifically these contaminants of concern are accumulating within ponds. This has implications for their impacts, since lead buried in the sediments of the pond within a solid compound is likely less bioaccessible to aquatic food webs than lead in its soluble form (McGeer et al. 2004). To accomplish this, we looked at the over 16,000 pond network of stormwater ponds across the state (MPCA 2025), focusing on the Twin Cities Metropolitan Area (a 7 county area containing Minneapolis and St. Paul, MN, USA). Similar to many older industrial cities, the Twin Cities Metro continues to grapple with persistent lead contamination and its associated health consequences (MN Department of Health 2023, Kemmerling et al. 2025). Research indicates that terrestrial lead distributions across the Twin Cities are associated with traffic volume and housing age (Mielke and Adams 1989, Shephard et al. 2022). The chemistry and ecology of stormwater ponds varies across the Twin Cities and it is unclear how this may impact lead sequestration and remobilization. To better mitigate the effects of lead contamination, we need understanding of the ecological processes that move lead through ecosystems. The urban lead cycle is inherently an ecological issue.

We explored three mutually non-exclusive hypotheses for the relationship between landscape variables and sediment lead accumulation, focusing on the impact of land use on soil contamination and potential transport pathways. (H1.1) Ponds that receive stormwater from older areas will have higher concentrations of lead due to a higher probability of lead paint participles in nearby soils. (H1.2) Ponds that receive stormwater from areas with high traffic volume will have high concentrations of lead due to past leaded gasoline inputs in nearby soils. (H1.3) Ponds that receive stormwater from areas with high impervious cover will have a lower concentration of lead due to reduced surface transport pathways.

We explored three hypotheses for the relationship between pond characteristics and lead dissolved within the water column, focusing on the role these features play in creating physical and chemical conditions that impact lead solubility. (H2.1) Ponds that are more resistant to mixing across the water column (i.e., highly stratified due to density differences between layers of water that vary in temperature and salinity) will have less lead dissolved within the surface water of the pond, since that water will rarely interact with lead contaminated sediments. Moreover, greater stratification is associated with anoxic conditions, especially at the bottom of these ponds, which could impact how much lead is dissolved based on whether lead is bound to organic material and/or sorbed to iron oxides. (H2.2) Ponds that have greater duckweed cover will have less lead dissolved within the surface water of the pond, since vegetative cover is also associated with the suppression of oxygen. (H2.3) Water with greater conductivity at both the surface and the bottom will have higher lead solubility due to increased capacity for cation exchange, since conductivity is a measure of chloride and calcium ions in the water.

We explored three hypotheses for where lead then accumulated within the pond structure itself. We hypothesized that (H3.1) lead will be at higher concentrations within the porewater and bottom water of ponds compared to the surface water, since that water will interact with the sediment bound lead at a greater rate. In addition, bottom water tends to be more anoxic, which could also impact how much lead is dissolved (similar to H2.1). Consequently, we also hypothesize that (H3.2) lead concentrations in vegetation at the surface of the pond, unrooted in sediment, should therefore correlate with surface water concentration. Moreover, we expect that as overall inputs have decreased, (H3.3) sediment lead concentrations will increase with increasing sediment depth.

## Methods

### Field collections and sample processing

Study sites were chosen from a database of ponds with water quality data from previous or ongoing studies and other supporting information that includes over 300 ponds across the Twin Cities Metro. Potential sites were narrowed down to where the predicted average building age was older than the 1970s or in areas with additional point sources of concern (i.e., near airports or where prior research had shown elevated stormwater lead (Finlay et al. 2024, Finlay et al. in preparation)). Sites were then prioritized to maximize variation in size, average annual traffic volume, and estimates of impervious surface cover within the data set.

Samples were collected between Summer 2023 and Winter 2025. One to three replicate sediment and water samples from 38 locations were collected, including 35 ponds and 3 urban lakes that receive stormwater inputs. Summer and fall samples (June – October) were collected from the surface of the pond via canoe to minimize mixing of pond layers while sampling. Winter (January – March) samples were collected from the surface of the pond through the ice via an ice auger. At each sampling location during the winter, two holes were drilled approximately 2 meters apart such that the collection of bottom water samples would not disturb the collection of chemical profiles of the pond. Sediment cores were then collected last at each sampling location. During the summer and fall, three to six samples of soil (10 cm cores) were collected around the ponds and analyzed for total lead using X-ray fluorescence (pXRF). When available, duckweed or filamentous algae were collected from the surface of the pond and then assessed for lead content by Inductively Coupled Plasma Optical Emission Spectroscopy (ICP-OES, iCap 7600 Duo Thermo Fisher Scientific) at the University of Minnesota Research Analytical Laboratory.

An HTH gravity coring device (Pylonex, Sweden) was used to collect sediment samples at up to three separate locations at each site, prioritizing locations within 3 m of an inlet, outlet, and toward the middle of the pond (if more than two samples were collected). Cores ranged in depth from only 10 cm to up to 50 cm in some locations. Sediment cores were sectioned every 10 cm and then sampled for total lead using X-ray fluorescence (pXRF). Bottom and surface water samples were collected using a peristaltic pump and Masterflex tubing. Only one surface water sample was collected at each location, since preliminary work indicated that surface water heavy metal concentrations were consistent throughout a single pond (Zweifel, unpublished data). Since bottom water concentrations varied across ponds, distinct bottom water samples were collected at each location where a sediment core was taken (1-3 per pond). Sediment porewaters were collected using 10 cm long MacroRhizon samplers (Rhizosphere Research Products, Wagingen, NL), which extract porewaters by filling evacuated glass vials and filtering to 0.15 µm. Water was analyzed for dissolved and total lead using an Inductively Coupled Mass Spectrometer (TQ-ICP-MS; iCAP TQ, Thermo Fisher Scientific).

### Estimating landscape variables

Parcel age was estimated as the average parcel construction year (MN Metropolitan Council MetroGIS), aggregated by the Minnesota Department of Natural Resources Level 09 autocatchments, natural landscape units which delineate catchments for lakes and provide a framework for organizing land use within watersheds (MN Department of Natural Resources Watershed Suite). Traffic volume was estimated as the annual average daily traffic of road segments within a 500 m radius of the pond (MN Department of Transportation Data Management System); we assumed that relative differences in traffic volume would be fairly consistent over time as much of the major road infrastructure in the state was built by the late 1960s. While this is an imperfect metric of historical traffic data, it should be indicative of traffic patterns when leaded fuels were prevalent. Previous work has found that 500 m is the buffer size corresponding to the mean drainage area for constructed stormwater basins with delineated catchments in the Twin Cities metro (n= 122 ponds; Janke et al. 2023). Mean impervious surface cover was calculated as the fraction of land covered by impervious cover surrounding the pond (MN Department of Natural Resources Watershed Suite). Distinct from parcel age, estimated basin age was calculated as the number of years since the pond was built or last dredged and the year of sampling (n = 32/38 sites). The size of the pond was determined using GIS to measure the surface area in acres. Mean depth of the pond was calculated as the average depth across the entire pond, either taken from prior data sets (Janke et al. 2023, Natarajan et al. 2025), or estimated from water profiles taken in the field (i.e., mean of depths measured). Soil lead was calculated by taking the mean of the estimates of soil lead from samples collected with 5 m of pond edge (see above).

### Estimating internal pond features

Mean dissolved oxygen (DO) was estimated by taking the average dissolved oxygen levels sampled vertically across the entire water column, collected at the same day and approximate location as the water sample. Similarly, relative thermal resistance to mixing (RTRM) for the whole water column was calculated for each specific location where a water column profile was collected within a pond. Since the ponds have high road salt inputs, we estimated water density using both temperature and conductivity (the latter to estimate salt concentration) (Janke et al. 2023). The pH of water samples was measured at the time of collection, and only for a subset of the ponds visited during the summer. Anoxic Factor (AF) was estimated as the fraction of the year during which an area equivalent to the pond surface area was exposed to anoxic water. In this case, AF was calculated from bathymetry and manual dissolved oxygen profiles and normalized by days in the year. AF was only estimated for a subset of ponds that were monitored across seasons (Natarajan et al. 2025). For the subset of ponds visited in the summer or fall, we also had an estimate of the percent of the pond visually covered in duckweed.

### Statistical analysis

We used generalized linear multilevel models in a Bayesian framework fitted with the R package brms to analyze predicted relationships (Bürkner 2018). We considered the magnitude of a fixed effect to be different from zero if the associated 95% credible interval of the posterior parameter estimate did not overlap with zero. For estimates of fixed effects in all models, we used weakly informative, regularizing priors centered on 0. For model selection, we used leave-one-out cross validation (LOO) to compare fit between models that represented the various hypotheses being tested for predicting each outcome variable. If multiple models were close in their LOO score, we added additional models that combined the variables from the top three lowest LOO scores and included those models in comparison. The posterior parameter values reported were from the model that had the lowest LOO criteria value across all models.

### Across pond variation: landscape level characteristics that impact lead accumulation in sediment

To evaluate differences in surface sediment lead concentrations (0-10 cm depth) based on landscape variables, we used a gaussian structure. We included varying intercepts for each pond as a random effect in our model. Moreover, only sites with estimated basin age were included in this analysis (n = 32), since dredging is an important mediator of sediment content. Thus, to control for the impact of dredging, basin age was included in all models compared. Parcel age, pond size, mean total impervious surface cover, and annual average daily traffic were all included as fixed predictors in the models compared during model selection. Since pond and lake catchments can differ quite dramatically in size, lakes were not included in this analysis. Initial model exploration indicated no evidence that depth or mean soil lead impacted sediment lead concentrations, so both were left out of the final model.

### Across pond variation: pond characteristics that impact dissolved lead within the water column

To evaluate differences in dissolved lead within surface water or bottom water based on internal pond features, we used a censored gamma structure for the response variable in this model. This is because water lead concentrations were left-censored, with over 15% of values of lead concentration below the limit of detection (0.2 ppb). The Bayesian approach to handling left-censored data can enhance analysis in complex datasets containing non-detects by modeling the non-detects as a distribution between zero and the limit of detection (Shoari and Dubé 2018). For the analysis of both surface and bottom water, summer and fall locations were analyzed separately from winter locations because frozen ponds can become inverse stratified (Ahmed et al. 2022) and seasonal variation in sediments inputs and water chemistry lead could change which parameters were most important for determining dissolved lead concentrations. In initial data visualization, mean DO across the entire water column was a better predictor of water lead levels than anoxic factor for the subset of ponds in which anoxic factor had been calculated. Thus, mean DO or surface/bottom DO was included as a predictor of dissolved oxygen levels in subsequent models. To assess the impact of pH on lead water concentrations, a separate analysis with the subset of data with known pH values at the time of collection was also performed. It confirmed that including pH did not improve model fit.

For analysis of surface water lead concentrations, mean pond depth, mean DO across the entire pond, surface DO, mean RTRM across the entire pond, surface water conductivity (as a proxy for salinity), surface water temperature and percent duckweed cover were all included as fixed predictors in model comparison. For analysis of bottom water lead concentrations, pond size, mean depth, mean DO across the water column at collection location, bottom water DO at collection location, RTRM at collection location, bottom water conductivity (as a proxy for salinity), bottom water temperature, and percent duckweed cover were all compared as fixed predictors. To control for variation in the amount of potentially water-soluble lead, sediment lead (top 10 cm) in the pond was included as a predictor in all models, either as mean across the pond (surface water analysis) or at the collection site (bottom water analysis). Since multiple bottom water samples were collected at each pond, we included varying intercepts for each pond as a random effect in this model.

For both analysis of surface and bottom water, all continuous variables were scaled to improve model convergence and make parameter interpretation easier. Additionally, to understand if similar patterns occurred even when ponds were frozen, a separate analysis was performed with the subset of samples collected in the winter (Appendix S1).

### Within pond variation: lead concentration spatial trends

To evaluate the relationship between lead water concentrations and location within the water column, we used a censored gamma structure. We included varying intercepts for each pond as a random effect in our model. Water column depth (i.e., surface water, bottom water, porewater) was included as a categorical predictor. To control for the amount of lead present in each pond and the different sizes of each pond, mean pond surface sediment lead (top 10 cm) and mean pond depth were included as continuous predictors. We used posterior odds ratios to interpret pairwise contrasts between the categorical variables in our model. We considered the differences in lead between water column depths to be different than zero if the associated 95% credible interval of the posterior odds ratio did not overlap 1.

To evaluate the relationship between vegetation lead concentrations and surface water concentration, we used a gamma structure to model the distribution of the response variable. Surface water lead concentration and vegetation type were included as predictors. To evaluate differences in sediment lead concentrations based on depth, we used a negative binomial structure due to the distribution of the response variable. We included a nested random effect structure, where there is a random intercept for each pond and within each pond, there is also a random intercept for each sampling location. Sediment depth was then included as a fixed predictor in the model.

During the development of the data analysis pipeline, Claude Sonnet 4.5 (Anthropic, 2025) was used to assist with troubleshooting error messages and improving visualization of conditional effects, including unscaling posterior parameter values for figures. The output was reviewed, tested, and substantially edited by the authors, who take full responsibility for the final code. The complete relevant code is available at https://github.com/leapollack/Pb_hotspots_ponds.

## Results

Across all collected water samples collected, 82% (31/38) had detectable levels (above 0.2 ppb for the sampling method) of lead in surface water samples and 81% (63/78) had detectable levels of lead in bottom water (Fig 1). All ponds had detectable levels of lead within sediment at all depths (range 8 – 363 ppm). Five ponds contained sediment sampled with lead levels above 200 ppm, the EPA recommended “screening level” for residential soils (EPA Soil Lead Guidelines 2024). Detailed posterior parameter summaries for the models referenced below can be found in Appendix S2.

**Fig 1.**
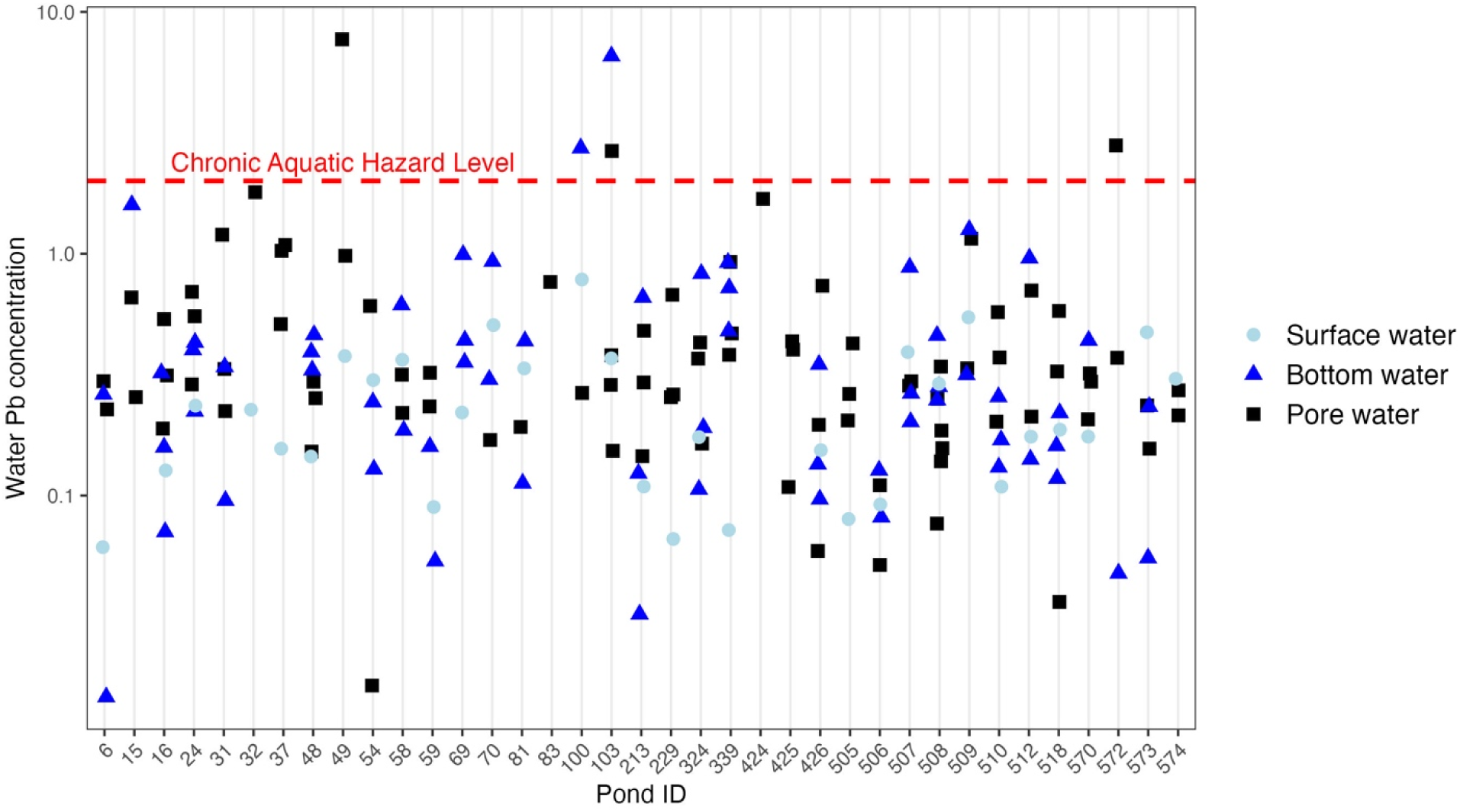
Raw data for lead concentrations in all water types collected for all non-frozen sites. Light blue circles are surface water, dark blue triangles are bottom water, and black squares are pore water. Note that values below the limit of detection are not depicted in this figure. The chronic aquatic hazard level for lead (EPA 1984) is indicated in red.

### Across pond variation: landscape level characteristics that impact lead accumulation in sediment

The best fit model predicting surface sediment lead concentrations included estimated parcel age (estimate: –2.31, CI: –3.58 – –1.07) and age of pond basin (estimate: –0.09, CI: –1.38 – 1.20) (Table 1, Fig 2). In support of H1.1, the best predictor of surface sediment lead concentrations was estimated parcel age if age of pond basin was known. The 95% CI for the posterior parameter estimates for parcel age did not overlap with zero, indicating that we are confident it is a strong predictor from this data set (Fig1). During model selection, including average annual traffic volume or mean impervious surface cover did not improve fit for the model of surface sediment lead concentrations, indicating that we did not find strong support for H1.2 or H1.3.

**Fig 2.**
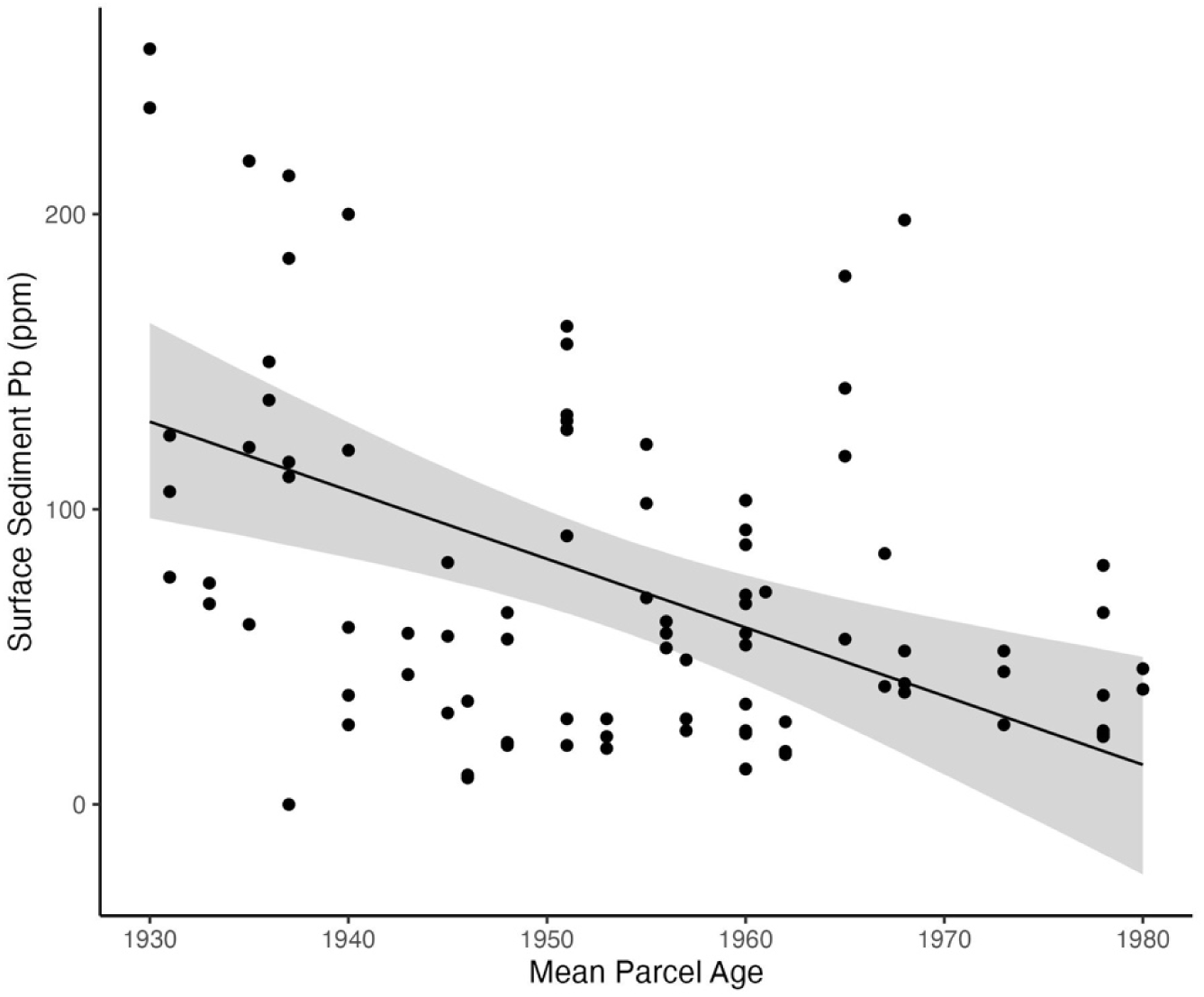
Raw data and conditional effects plots of mean parcel on lead concentrations in the top 10 cm of pond sediment. The conditional effects show the global effects of mean parcel age on sediment lead concentrations using expected values from the posterior parameter distribution of the best fit model. The lines represent the median effects, while the bands show the 95% credible intervals for the conditional effects. Black dots represent raw data values (i.e., individual sediment samples, n = 1-3 samples for 32 ponds).

Table 1.

Table 1.
| Model structure | ELPD<br>Difference | SE<br>Difference |
| --- | --- | --- |
| sediment Pb ~ parcel age + basin age + (1\| pond) | 0.0 | 0.0 |
| sediment Pb ~ parcel age + basin age + size + (1\| pond) | -0.1 | 0.4 |
| sediment Pb ~ parcel age + basin age + impervious surface cover + (1\| pond) | -0.1 | 0.4 |
| sediment Pb ~ parcel age + basin age + traffic volume + (1\| pond) | -0.3 | 0.6 |
| sediment Pb ~ traffic volume + basin age + size + (1\| pond) | -1.2 | 1.4 |
| sediment Pb ~ basin age + size + (1\| pond) | -1.5 | 1.4 |
| sediment Pb ~ impervious surface cover + basin age + size + (1\| pond) | -1.6 | 1.5 |

*ELPD Difference is the difference in the estimated expected log predictive density between each* *model and the best-performing model. The model with the highest ELPD is in the first row with* *an ELPD difference of 0. SE Difference is the standard error in the ELPD Difference.*

### Across pond variation: pond characteristics that impact dissolved lead within the water column

The best fit model predicting surface water lead concentrations included RTRM as a fixed effect (Table 2). In support of H2.1, RTRM was the best predictor of surface water lead concentrations (estimate: 0.55, CI: 0.01 – 1.20), but in the opposite direction than expected: ponds that were more resistant to mixing had higher lead within the surface water (Fig 3a). It is important to note that the uncertainty in the model’s predictions increases quickly at higher values. In other words, the predictions at high lead concentration in water are poor. This trend, that ponds with higher RTRM had higher lead in the surface water, was not supported in the smaller subset of frozen ponds (Appendix S1). Notably, surface sediment lead (top 10 cm) was not a strong predictor of surface water lead concentrations, as its 95% credible interval overlapped with zero (estimate: 0.14, CI: –0.28 – 0.69).

**Fig 3.**
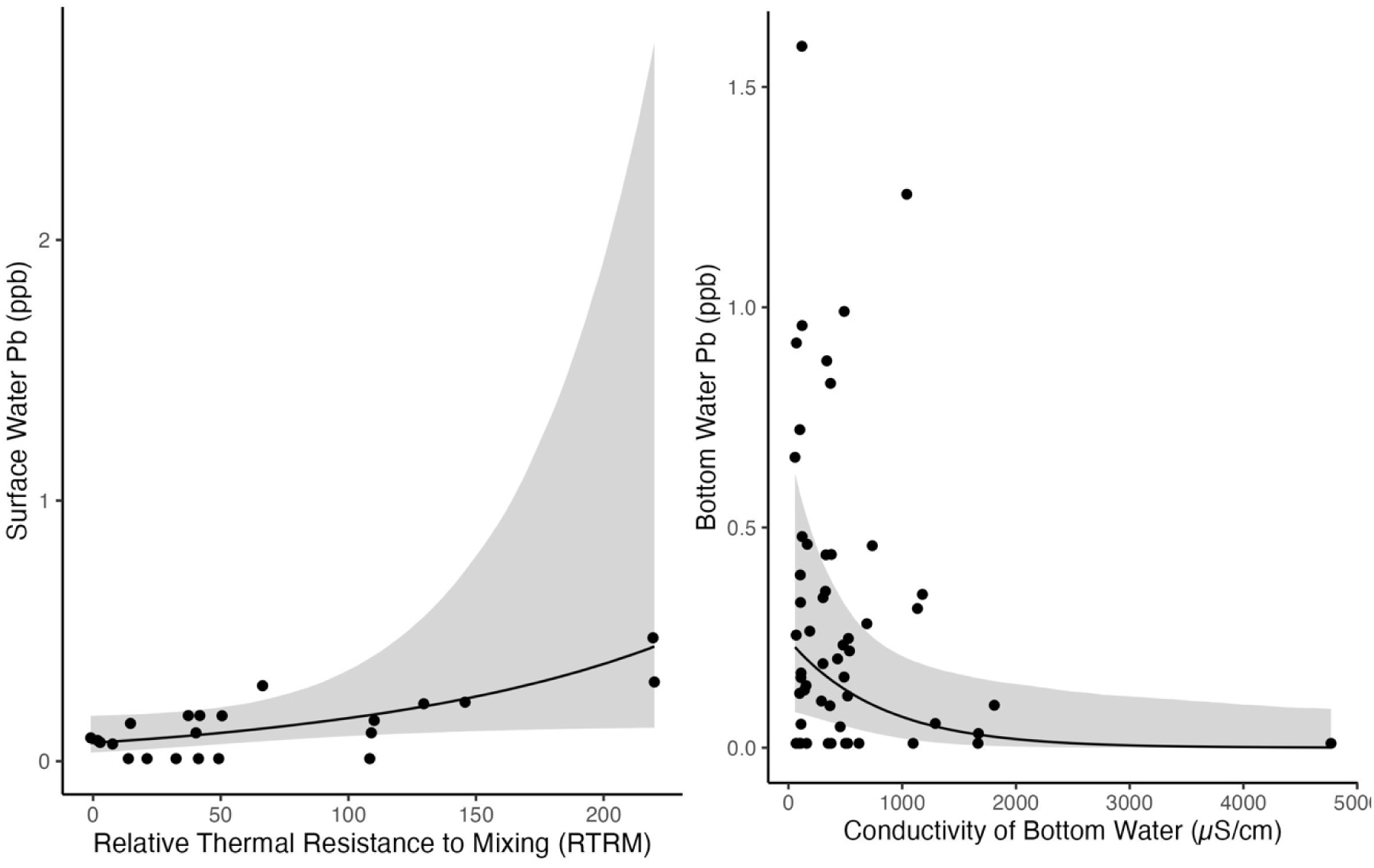
Raw data and conditional effects plots of (a) relative thermal resistance to mixing (RTRM) on surface water lead concentrations and (b) conductivity on bottom water lead concentrations. The conditional effects show the global effects of these proxies for pond stratification on water lead concentrations using expected values from the posterior parameter distribution of the best fit models for each water depth respectively. The lines represent the median effects, while the bands show the 95% credible intervals. Black dots represent raw data values.

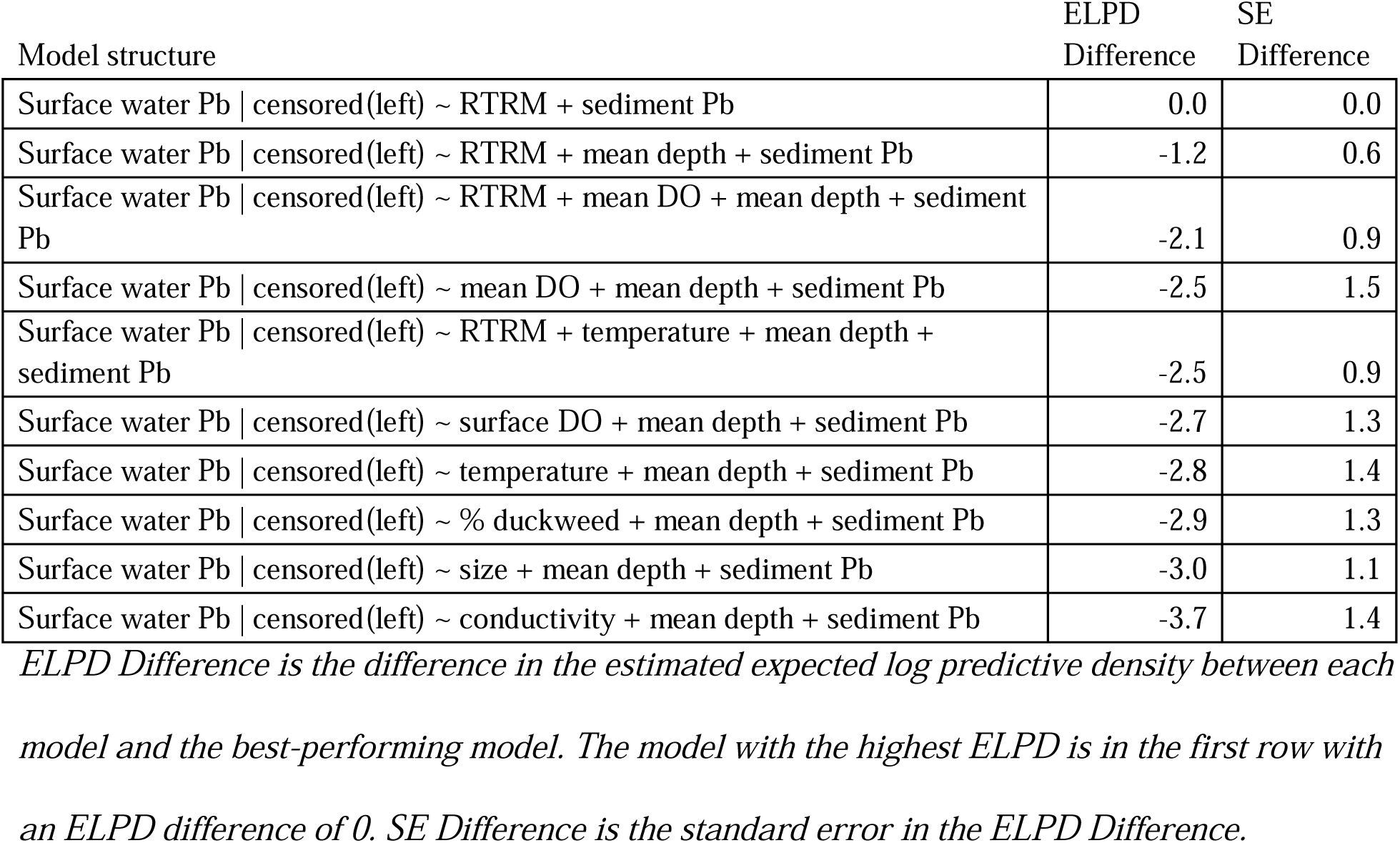
Table 2.

Table 2.
| Model structure | ELPD<br>Difference | SE<br>Difference |
| --- | --- | --- |
| Surface water Pb \| censored(left) ~ RTRM + sediment Pb | 0.0 | 0.0 |
| Surface water Pb \| censored(left) ~ RTRM + mean depth + sediment Pb | -1.2 | 0.6 |
| Surface water Pb \| censored(left) ~ RTRM + mean DO + mean depth + sediment Pb | -2.1 | 0.9 |
| Surface water Pb \| censored(left) ~ mean DO + mean depth + sediment Pb | -2.5 | 1.5 |
| Surface water Pb \| censored(left) ~ RTRM + temperature + mean depth + sediment Pb | -2.5 | 0.9 |
| Surface water Pb \| censored(left) ~ surface DO + mean depth + sediment Pb | -2.7 | 1.3 |
| Surface water Pb \| censored(left) ~ temperature + mean depth + sediment Pb | -2.8 | 1.4 |
| Surface water Pb \| censored(left) ~ % duckweed + mean depth + sediment Pb | -2.9 | 1.3 |
| Surface water Pb \| censored(left) ~ size + mean depth + sediment Pb | -3.0 | 1.1 |
| Surface water Pb \| censored(left) ~ conductivity + mean depth + sediment Pb | -3.7 | 1.4 |

*ELPD Difference is the difference in the estimated expected log predictive density between each* *model and the best-performing model. The model with the highest ELPD is in the first row with* *an ELPD difference of 0. SE Difference is the standard error in the ELPD Difference.*

The best fit model predicting bottom water lead concentrations included percent duckweed cover (estimate: –0.90, CI: –1.98 – 0.12) and bottom water conductivity as fixed effects (estimate: –0.91, CI: –1.71 – –0.09) (Table 3). In support of H2.2, percent duckweed cover was the best fit predictor of lead in the water column, but in the bottom, not surface water. It is important to note that the 95% CI for the posterior parameter estimate of percent duckweed cover slightly overlaps with zero, indicating we are less confident in this trend than the estimated effect of conductivity. Moreover, neither variable was in the best fit model describing trends observed in the winter ponds (Appendix S1); however, duckweed cover could not be included as a variable for this data set because the ponds were frozen over. In support of H2.3, conductivity was an important predictor of dissolved lead, but not in the direction expected (Fig 3b): ponds with higher conductivity were predicted to have lower bottom water lead concentrations, but the pattern was opposite. Similar to surface water, sediment lead was not a strong predictor of bottom water lead concentrations either, as its 95% credible interval overlapped with zero (estimate: 0.08, CI: –0.50 – 0.70).

Table 3.

Table 3.
| Model structure | ELPD<br>Difference | SE<br>Difference |
| --- | --- | --- |
| Bottom water Pb \| censored(left) ~ conductivity + % duckweed + sediment Pb + (1\| pond) | 0.0 | 0.0 |
| Bottom water Pb \| censored(left) ~ conductivity + % duckweed + mean depth + sediment Pb + (1\| pond) | -0.4 | 0.4 |
| Bottom water Pb \| censored(left) ~ temperature + mean depth + sediment Pb + (1\| pond) | -1.2 | 1.1 |
| Bottom water Pb \| censored(left) ~ conductivity + mean depth + sediment Pb + (1\| pond) | -1.2 | 0.6 |
| Bottom water Pb \| censored(left) ~ conductivity + temperature + mean depth + sediment Pb + (1\| pond) | -1.6 | 0.8 |
| Bottom water Pb \| censored(left) ~ conductivity + RTRM + mean depth + sediment Pb + (1\| pond) | -2.2 | 0.8 |
| Bottom water Pb \| censored(left) ~ RTRM + mean depth + sediment Pb + (1\| pond) | -2.7 | 1.0 |
| Bottom water Pb \| censored(left) ~ % duckweed + mean depth + sediment Pb + (1\| pond) | -2.8 | 1.7 |
| Bottom water Pb \| censored(left) ~ size + mean depth + sediment Pb + (1\| pond) | -3.1 | 1.7 |
| Bottom water Pb \| censored(left) ~ mean DO + mean depth + sediment Pb + (1\| pond) | -4.3 | 1.5 |
| Bottom water Pb \| censored(left) ~ bottom DO + mean depth + sediment Pb + (1\| pond) | -5.0 | 1.8 |

*ELPD Difference is the difference in the estimated expected log predictive density between each* *model and the best-performing model. The model with the highest ELPD is in the first row with* *an ELPD difference of 0. SE Difference is the standard error in the ELPD Difference.*

We did not find support for the hypothesis that dissolved oxygen was a good predictor of water lead levels (H2.3), as it did not improve model fit for predicting lead levels in surface or bottom water.

### Within pond variation: lead concentration spatial trends

We found that, on average, surface water had lower lead levels than bottom water (median odds ratio estimate bottom – surface: 1.89, CI: 1.04 – 2.90) and porewater (median odds ratio estimate pore – surface: 2.55, CI: 1.40 – 3.90), in support of H3.1. We did not find a difference in lead concentrations between bottom water and the top 10 cm of porewater (median odds ratio estimate bottom – pore: 0.72, CI: 0.46 – 1.07). We failed to find support for H3.2, as we did not observe that vegetation lead concentrations correlated with surface water values (estimate 0.30, CI: –4.61 – 5.86). Importantly, we found that sediment depth matters. Lead concentrations increased within deeper sediment (estimate 0.02, CI: 0.01 – 0.02), providing support for H3.3 (Fig 4).

**Fig 4.**
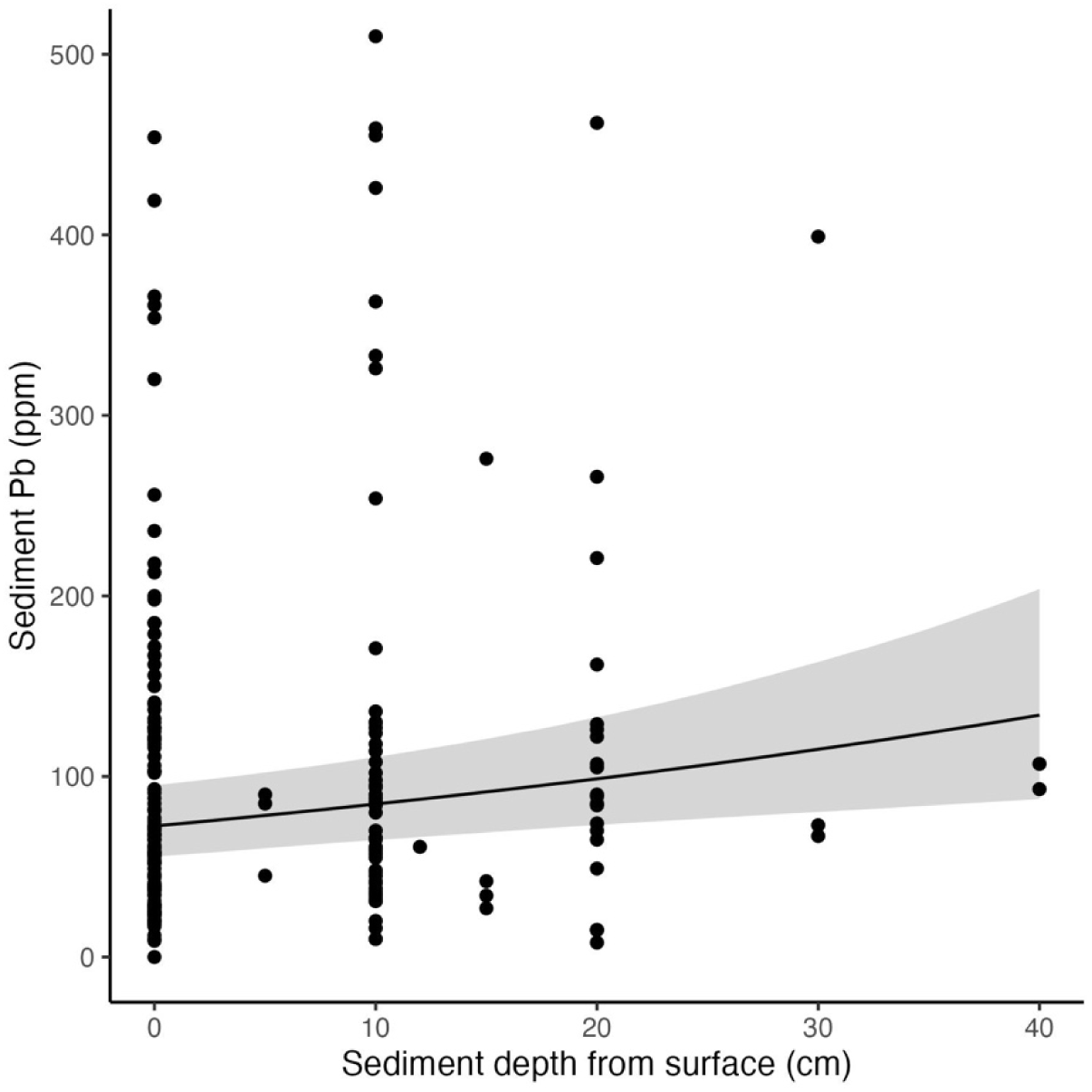
Raw data and conditional effects plots of sediment depth on sediment lead concentrations. The conditional effects show the global effects of sediment depth on lead concentrations using expected values from the posterior parameter distribution of model. The lines represent the median effects, while the bands show the 95% credible intervals. Black dots represent raw data values. This plot is a magnified view of the raw data with sediment lead values less than 500 ppm to show the conditional effects pattern more clearly.

## Discussion

The age of surrounding infrastructure, when age of the basin is known, stands out as the most robust predictor of sediment lead concentrations in urban stormwater ponds within this study. While multiple environmental and design-related factors can influence heavy metal accumulation in stormwater ponds (Egemose et al. 2015), these findings emphasize the dominant role of legacy materials and infrastructure wear, which likely contribute persistent sources of lead to urban runoff over time. Indeed, lead oxide, indicative of lead paint pigment sources, has been found in high proportion compared to other lead containing compounds within urban soils across the Twin Cities Metro (Zweifel et al., unpublished data). Moreover, our findings complement studies in other urban regions of the U.S., which have shown a positive relationship between distance to aging infrastructure and contaminated soils (Filippelli et al. 2018, Wade et al. 2021). Similarly, research on the stormwater inputs within the Twin Cities Metro has found a pattern of higher lead concentrations within watersheds with older infrastructure (Finlay et al. 2024). This relationship underscores how historical urban development patterns and aging building materials continue to shape contemporary aquatic pollution, highlighting the importance of considering infrastructure timelines when assessing and managing sediment contamination in urban water bodies.

Physical and chemical properties of the pond had a stronger impact on dissolved lead levels within the water column than the lead concentrations within pond sediment. Surprisingly, ponds with hallmarks of higher stratification (i.e., higher RTRM and greater duckweed cover), tended to have higher lead levels in their surface water and lower lead levels in their bottom water. This pattern was both unexpected and not strongly supported by model outcomes for bottom water. Thus, we view this finding as hinting at a possible trend more than explaining clear differences between lead levels across ponds. While unexpected, this trend could possibly be explained by photochemical reactions (e.g., reduction of iron oxides or organic material, Kopáček et al. 2006, Shiller et al. 2006, Kim et al. 2019, Lueder et al. 2020) that release lead at the surface and stratification that then slows mixing of the dissolved lead across the water column. However, it is important to note that evidence from daily monitored ponds in the area suggests that even the most intensely-stratified ponds mix partially on a nearly daily basis, such that surface water is likely interacting with the lower water column (Janke 2023). While stratification would promote anoxic conditions in the bottom water, causing reduction of iron oxides and release of lead into the porewater and bottom waters, the strong affinity of lead to particulate organic matter would mean dissolved lead should not shift dramatically across redox transitions (Dewey et al. 2021). This might explain why we did not find that low oxygen levels in the bottom water were associated with an increase in dissolved lead.

Related to these trends, the strongest predictor of water lead levels was conductivity of bottom water. That is, more saline water had lower lead levels at the bottom of the pond, which is the opposite pattern expected based on the chemical properties of chloride and calcium rich water (Acosta et al. 2011). This suggests that runoff composition could be an important factor determining the patterns we are observing, with inputs that are both lower in salinity and higher in lead sitting on the surface of the ponds (for example, perhaps residential areas with high lead paint but relatively lower road salt inputs). Ultimately, these unexpected trends are quite small, mostly describing differences less than 1 ppb in dissolved lead levels across ponds. For context, the chronic aquatic hazard level is 2 ppb (U.S. EPA 1984) and the action level for drinking water is 10 ppb (U.S. EPA 2025). Thus, although notable differences in dissolved lead concentrations were observed among the ponds, the complexity of site-specific factors likely prevents us from making broad generalizations that explain the subtle variations in dissolved lead levels between ponds.

Overall, our findings suggest that stormwater ponds are effective at capturing lead from urban runoff. For one thing, lead in pond sediments was not a strong predictor of dissolved lead (i.e., more bioavailable lead) within the water column. While we looked at ponds with a wide range of sediment lead concentrations, overall lead levels were low within the water column regardless of sediment levels. Moreover, lead levels in surface water were on average lower than concentrations at the bottom of the water column. In addition, we did not observe that water lead levels were a strong predictor of floating duckweed and algae lead levels (although this was a small sample size). This could suggest that the small differences in the amounts of potentially bioavailable lead did not necessarily cause differences in stoichiometric uptake. This is not surprising, as most plant species reduce lead absorption and often sequester it in their root tissues as a defense mechanism (reviewed in Pourrut et al. 2011). Moreover, lead levels tended to increase in deeper sediments, suggesting the burial of past higher inputs over time. Thus, not only are decreases in lead sources reflected in the sediment record (similar to Hill et al. 2023), but past leaded inputs are effectively being sequestered deep within pond sediment where it is less likely to interact with local pond food webs.

This study has several limitations that should be considered when interpreting these results. For one, many of the ponds studied had one to multiple intake pipes buried underground, likely affecting the relative importance of surface transport pathways for inputs across ponds. Moreover, where samples were taken in respect to these inlet pipes might affect heavy metal concentrations, since distance to inlets can impact heavy metal concentration variation across the pond floor (Crawford et al. 2010, Egemose et al. 2015). While we attempted to collect 2 – 3 sediment samples per pond, focusing on having a sampling location near inlet, outlets, and the center of the pond, we might have missed clumps of spatial variation in heavy metals across pond sediments. Second, we did not incorporate seasonal variation within our study due to time constraints. Instead, we removed frozen ponds from the main data set when analyzing lead levels in the water column due to known seasonal differences across ponds in water chemistry.

Intriguingly, we did not find that the same predictor variables were important in the small subset of frozen ponds we analyzed (Appendix S1). This suggests that there are a lot of seasonal variations both in sediment inputs (e.g., storm events, spring melt) and water chemistry (e.g., changes in stratification, anoxia, salt levels) that could be playing an important role in describing differences in water lead levels. Lastly, the analysis for the relationship between water lead levels and pond vegetation was quite small and focused on plants not rooted in the sediment, where most of the lead within these ponds was concentrated. Thus, although we did not find a positive relationship between water lead levels and floating vegetation lead concentrations, we might still observe differences in bioaccumulation of lead for plants that are rooted in leaded sediments.

### Ecological implications

The variation observed in lead concentrations within stormwater ponds has important ecological implications. Our findings suggest that pond sediments act as sinks in the urban lead cycle, accumulating contaminants over time. However, ponds could also be potential sources of lead if they release lead under changing environmental conditions such as shifts in pH, increased salinity, or disturbance events. The finding that most lead within ponds is found within the sediment suggests that benthic invertebrates and other bottom-feeding species (e.g., snails, crayfish, minnows) may experience greater exposure risks than pelagic species that utilize these ponds. Indeed, stormwater ponds in Minnesota can support a variety of fish species, including fathead minnows, bullheads, goldfish, carp, and green sunfish (McComas and Stuckert. 2008, 2011). Meanwhile, relatively small amounts of lead in the water column indicates limited immediate bioavailability to pelagic organisms (e.g., daphnia, mosquitofish), as underscored by our findings on floating pond vegetation. While some bottom and pore water samples within our sample set exceeded the chronic aquatic hazard level for lead (Fig 1), these were relatively rare (i.e., only three bottom water samples had lead concentrations above 2 ppb). Interestingly, while we did not find that water lead levels predicted vegetation lead levels, we did observe a few duckweed and algae samples that were relatively high in lead. This might be an anomaly or perhaps hints at a process we have yet to understand given the complex differences in chemistry and nutrient cycling across ponds. The strong differences in sediment lead concentrations across stormwater ponds, contrasted with relatively minimal differences in pond water lead levels, underscores that landscape-level spatial variation might be more important than individual pond characteristics for predicting aquatic lead hotspots. This suggests that broader environmental and land use patterns, such as urban infrastructure age, drive the heterogeneous distribution of lead contamination more than specific pond design or management features.

### Management implications

The finding that the age of surrounding infrastructure is a stronger predictor of sediment lead levels compared to other landscape features carries significant management implications. It highlights the necessity of targeting legacy sources of lead contamination linked to older urban development, such as aging building materials, to effectively reduce sediment lead hotspots in stormwater ponds. Since variations in water lead concentrations were minimal, management efforts should prioritize controlling sediment-bound lead, which represents a longer-term reservoir of contamination and potential secondary source under changing environmental conditions. Therefore, management strategies should consider risks associated with sediment disturbance. For example, if stormwater ponds were to dry out, the lead previously sequestered in sediments could become mobile again in the form of contaminated soil and dust, posing renewed ecological and human health risks. If dredging is planned, it is critical to estimate potential lead content in the dredged material accurately to inform disposal options and costs, as contaminated sediments require special handling (MPCA 2024) and can significantly increase project expenses. Additionally, sediment resuspension, such as might occur if fish species like koi or carp are introduced and disturb the pond bottom, could mobilize sediment-bound lead into the water column, increasing exposure risk to aquatic organisms. Biochar amendments may offer a potential method for stabilizing lead in pond sediments, but it remains unclear whether its effectiveness varies with pond chemistry (Cairns et al. 2022, Li et al. 2022, Thaçi et al. 2024). Collectively, these considerations highlight the need for integrated management approaches that address both the legacy contamination linked to aging infrastructure and the dynamic physical processes that influence lead availability in stormwater pond ecosystems.

## Acknowledgements

Thank you to Nic Jelinski, Nora Pearson, Catherine Polik, Tim Mitchell, Luis Santiago-Rosario, and Lindsey Kemmerling for advice while designing this experiment. Thank you to Madeline Taylor, Alexandria Wenger, Jackson Peterson, Ian Coffman, Mitchell Pariseau, Claire Bass, Lay Lay, Caroline Castellon, Jenna Duncan and Jake Wiebe for assisting in the field and laboratory. We thank the City of Minneapolis, Minneapolis Parks and Recreation, Minnesota Department of Transportation, Saint Paul Parks and Recreation, the City of Roseville, the City of Bloomington, Falcon Heights, Capitol Region Watershed District, and the University of Minnesota for providing access to stormwater ponds for research sampling. We also acknowledge the use of Claude Sonnet 4.5 (Anthropic, 2025), a generative artificial intelligence tool, for assistance in refining code in R for data analysis and figure generation. All analytical decisions and interpretations remain solely those of the research team.

