## Appendix S1 for "Predicting lead hotspots in urban stormwater ponds across the Twin Cities Metro"

**Appendix S1. Analysis predicting dissolved lead in the water of frozen ponds**

To understand if similar patterns to unfrozen ponds occurred even when ponds were frozen, a separate analysis was performed with all water samples collected in in the winter under the ice.

*Methods:*

The statistical analysis approach was the same as described in the main manuscript. That is, a Bayesian framework in which multi-level models were fitted with the R package brms and inference was made based on the 95% credible interval of the posterior parameter estimates. To control for variation in the amount of potentially water-soluble lead, sediment lead (top 10 cm) in the pond was included as a predictor in all models, either as mean across the pond (surface water analysis) or at the collection site (bottom water analysis). Since multiple bottom water samples were collected at each pond, we included varying intercepts for each pond as a random effect in this model. To evaluate differences in dissolved lead in the frozen ponds we used a gamma structure for the response variable in these models, since all values for this subset of the data had bottom water lead values above the limit of detection.

*Surface water:* For analysis of surface water lead concentrations, mean pond depth, mean DO across the entire pond, mean RTRM across the entire pond, surface water conductivity (as a proxy for salinity), surface water temperature and percent duckweed cover were all included as fixed predictors in model comparison. Since surface DO was not collected at all sites in the winter due to temperature constraints on the probe, a separate analysis with the subset of data with known surface DO values at the time of collection was also performed. It confirmed that including surface DO did not improve model fit compared to mean pond DO.

*Bottom water*: For analysis of bottom water lead concentrations, pond size, mean depth, mean DO across the water column at collection location, bottom water DO at collection location, RTRM at collection location, bottom water conductivity (as a proxy for salinity), and bottom water temperature.

*Results and Discussion:*

The best fit model predicting surface water lead concentrations for the frozen ponds included temperature (estimate: 0.33, CI: 0.00 – 0.70) as a fixed effect (Table A1). The best fit model predicting bottom water lead concentrations for the frozen ponds included temperature (estimate: -1.01, CI: -2.00 – 0.20) and RTRM as fixed effects (estimate: 0.66, CI: -0.09 – 1.34) (Table A2). It is important to note that the 95% CI for the posterior parameter estimates for all variables cover overlaps with zero, indicating we are not as confident in this trend compared to other findings in this study.

Intriguingly, these are different than the best fit models from the unfrozen data set and both include temperature as an important predictor for differences in lead content across the frozen ponds. However, it is important to note that these are quite small datasets (n = 8 ponds), thus it is not surprising it is hard to make strong inference about which variables are making predictions across these ponds.

| Model structure | ELPD Difference | SE Difference |
| --- | --- | --- |
| Surface water Pb ~ temperature + mean depth + sediment Pb | 0.0 | 0.0 |
| Surface water Pb ~ temperature + mean DO + mean depth + sediment Pb | -2.3 | 1.2 |
| Surface water Pb ~ temperature + RTRM + mean depth + sediment Pb | -2.3 | 1.3 |
| Surface water Pb ~ mean DO + mean depth + sediment Pb | -2.6 | 1.8 |
| Surface water Pb ~ RTRM + mean depth + sediment Pb | -3.2 | 1.5 |
| Surface water Pb ~ conductivity + mean depth + sediment Pb | -3.7 | 1.2 |

*Table S1. ELPD Difference is the difference in the estimated expected log predictive density between each model and the best-performing model. The model with the highest ELPD is in the first row with an ELPD difference of 0. SE Difference is the standard error in the ELPD Difference.*

| Model structure | ELPD Difference | SE Difference |
| --- | --- | --- |
| Bottom water Pb ~ temperature + RTRM + mean depth + sediment Pb + (1\| pond) | 0.0 | 0.0 |
| Bottom water Pb ~ temperature + mean depth + sediment Pb + (1\| pond) | -3.4 | 1.6 |
| Bottom water Pb ~ temperature + conductivity + mean depth + sediment Pb + (1\| pond) | -3.6 | 1.4 |
| Bottom water Pb ~ RTRM + mean depth + sediment Pb + (1\| pond) | -4.4 | 1.1 |
| Bottom water Pb ~conductivity + mean depth + sediment Pb + (1\| pond) | -4.8 | 1.0 |
| Bottom water Pb ~ mean DO + mean depth + sediment Pb + (1\| pond) | -5.2 | 1.5 |
| Bottom water Pb ~conductivity + RTRM + mean depth + sediment Pb + (1\| pond) | -5.2 | 1.2 |
| Bottom water Pb ~ bottom DO + mean depth + sediment Pb + (1\| pond) | -5.5 | 1.3 |

*Table S2. ELPD Difference is the difference in the estimated expected log predictive density between each model and the best-performing model. The model with the highest ELPD is in the first row with an ELPD difference of 0. SE Difference is the standard error in the ELPD Difference.*
