## Appendix S2 for "Predicting lead hotspots in urban stormwater ponds across the Twin Cities Metro"

**Appendix S2. Model structure and posterior parameter estimates for models**

| **Model Structure** | **Posterior parameter estimates for fixed effects** | | | |
| --- | --- | --- | --- | --- |
|  | Parameter | Estimate | 2.5% CI | 97.5% CI |
| **Surface sediment Pb ~** parcel age + basin age + (1\| pond) | intercept | 4592.00 | 2165.34 | 7067.55 |
|  | basin age | -0.09 | -1.38 | 1.20 |
|  | parcel age | -2.31 | -3.58 | -1.07 |
|  | sigma | 34.52 | 28.60 | 42.09 |
| **Surface water Pb \| censored(left)** ~ RTRM + sediment Pb | intercept | -2.06 | -2.57 | -1.45 |
|  | sediment Pb | 0.14 | -0.28 | 0.69 |
|  | RTRM | 0.55 | 0.01 | 1.20 |
|  | shape | 0.68 | 0.32 | 1.20 |
| **Bottom water Pb \| censored(left)** ~ conductivity + % duckweed + sediment Pb + (1\| pond) | intercept | -2.13 | -3.12 | -1.22 |
|  | % duckweed | -0.90 | -1.98 | 0.12 |
|  | conductivity | -0.91 | -1.71 | -0.09 |
|  | sediment Pb | 0.07 | -0.49 | 0.68 |
|  | shape | 1.10 | 0.59 | 1.75 |
| **Water Pb \| censored(left) ~** water column depth + sediment Pb + pond depth + (1\| pond) | intercept | -0.97 | -1.46 | -0.49 |
|  | water column BW vs PW | 0.30 | -0.10 | 0.70 |
|  | water column BW vs SW | -0.63 | -1.12 | -0.12 |
|  | sediment Pb | 0.00 | 0.00 | 0.00 |
|  | pond depth | -0.03 | -0.12 | 0.06 |
|  | shape | 0.64 | 0.52 | 0.77 |
| **Vegetation Pb ~** surface water Pb + vegetation type | intercept | 1.79 | -0.03 | 3.69 |
|  | surface water Pb | 0.30 | -4.61 | 5.86 |
|  | vegetation type | 0.03 | -1.49 | 1.42 |
|  | shape | 0.83 | 0.37 | 1.51 |
| **Sediment Pb ~** depth + (1\| pond/location) | intercept | 4.29 | 4.01 | 4.56 |
|  | sediment depth | 0.02 | 0.01 | 0.02 |
|  | shape | 4.08 | 3.13 | 5.21 |
| *Note continuous variables for models of surface water and bottom water lead concentrations were scaled to improve model convergence and make parameter interpretation easier.* | | | | |
